# Safeguarding the genetic integrity of native pollinators requires stronger regulations on commercial lines

**DOI:** 10.1101/783878

**Authors:** Ignasi Bartomeus, Francisco P. Molina, Amparo Hidalgo-Galiana, Joaquín Ortego

**Affiliations:** Department of Integrative Ecology, Estación Biológica de Doñana, EBD-CSIC, Avda. Américo Vespucio 26, Seville E-41092, Spain

**Keywords:** bumblebees, hybridization, managed pollinators, subspecies.

## Abstract

Every year more than one million commercial bumblebee colonies are deployed in greenhouses worldwide for its pollination services to several commercially important crops such as tomato and different species of berries. While commercial pollinators have been an enormous benefit for the production of essential food crops and for achieving higher yields and better fruit quality at a low cost, their use is emerging also as an important threat to wild pollinators. Commercial pollinators have been linked to pathogen spillover to wild species, and its introduction outside its native area have had devastating effects on native pollinator populations. However, a more pervasive, but underappreciated threat is their potential impact on the genetic integrity of native pollinators. Here, we show clear evidence of generalized hybridization between native and introduced commercial bumblebee lineages in southern Spain. The signal of genetic introgression is widespread and already expands up to 60 km from main commercial bumblebee release areas. As pollination services demand is predicted to increase in the coming years, only a more restrictive regulation of commercial lines could mitigate their negative impacts on the genetic integrity of native pollinators and prevent the disruption of local adaptations.

## 1. INTRODUCTION

In 1987 commercial rearing of bumblebees started in the Netherlands for the pollination of tomato crops. Nowadays, more than 30 commercial producers worldwide supply pollination services in more than 60 countries (Velthuis & van Doorn, 2006). While five species of bumblebees are reared commercially, most of the market is dominated by two species: *Bombus terrestris* and *B. impatiens*. *Bombus terrestris* colonies have been used for commercial pollination not only in its Eurasian native range but also in East Asia (Japan, South Korea, China), South America (Chile) and New Zealand, and the eastern North American *B. impatiens* has been used in western North America and Mexico. As each bumblebee colony can produce over 200 queens, it is not surprising that commercial species have escaped into the wild and established naturalized populations in the introduced areas. The consequences for native pollinators, including direct competition and the spread of pathogens (Colla, Otterstatter, Gegear, & Thomson, 2006), have been in some cases devastating (e.g. the decline of *B. dahlbomii* in Chile; Morales, Arbetman, Cameron, & Aizen, 2013) and most countries, but not all, regulate nowadays the import of exotic species (Aizen et al., 2019).

The trade of bumblebees within its natural area of distribution contains a more silent threat. *Bombus terrestris* is a widespread species divided into nine well-defined subspecies with contrasting coloration patterns and local geographical adaptations (Rasmont, Coppee, Michez, & De Meulemeester, 2008). For example, while northern European subspecies hibernate, awakening from diapause in spring, southern subspecies aestivate and start their cycle in autumn. However, commercial colonies of some of the subspecies of *B. terrestris* have been widely used outside their natural distribution area. Several subspecies of *B. terrestris* were initially used in the early years of commercial rearing, but from the commercial point of view, *B. terrestris dalmatinus* proved to have superior characteristics and is the most common sold subspecies nowadays (Velthuis & van Doorn, 2006). Selling companies often argue that queen production of commercial colonies, escape from greenhouse conditions, and survival in the wild is unlikely. This view has resulted in no measures taken in most countries to regulate subspecies trade within Europe. In contrast, evidence is piling up that both male and queen production of commercial colonies are high, the produced queens can survive in the wild (Owen, Bale, & Hayward, 2016) and mating is not only happening among subspecies (Ings, Raine, & Chittka, 2005), but also among related species (Kondo et al., 2009). The genetic risks associated with releases of commercial species are largely neglected in conservation plans (Laikre, Schwartz, Waples, Ryman, & Ge, 2010), however, genetic pollution can lead to the breakdown of coadapted genes complexes, erode local adaptation processes, and reduce the ability of populations to deal with different components of global change. Unfortunately, economic interest usually dominates decision-making and in the absence of solid evidence of genetic pollution from commercial pollinators, only a few countries (see below) have regulated the genetic lines that can be commercially used.

Spain is one of the main vegetable producers in Europe and several crops, mainly tomato and different species of berries, use commercial bumblebees to supplement pollination. Commercial bumblebees are used in Spain since 1992 and albeit actual commercial species are not necessarily pure lines, most sold bumblebees probably belong to the subspecies *B. t. dalmatinus* or *B. t. terrestris*. However, in the Iberian Peninsula the native subspecies is *B. t. lusitanicus*, a taxon characterized by its distinctive legs with reddish setae (Rasmont, Coppee, Michez, & De Meulemeester, 2008). Recent studies show that the commercial subspecies actively forage in natural areas and can produce viable queens (Trillo, Brown, & Vilà, 2019). Hence, we set up a sampling and genotyping protocol to evaluate the presence and prevalence of hybridization between commercial individuals and the native subspecies.

## 2. METHODS

### 2.1. Sampling

During 2017 and 2018, we collected via sweep-netting a total of 66 free-foraging individuals of *B. terrestris* from 28 sampling sites located at different distances from main greenhouse areas in southwest Andalusia, Spain (Figure 1; Table S1). Additionally, we sampled four individuals from purchased commercial colonies of the two main companies operating in the region (Agrobío S.L. and Koppert España S.L.; Table S1). We placed all sampled specimens in vials with 4 mL of ethanol 96% and stored them at −20° C until needed for phenotypic and genomic analyses.

**FIGURE 1.**
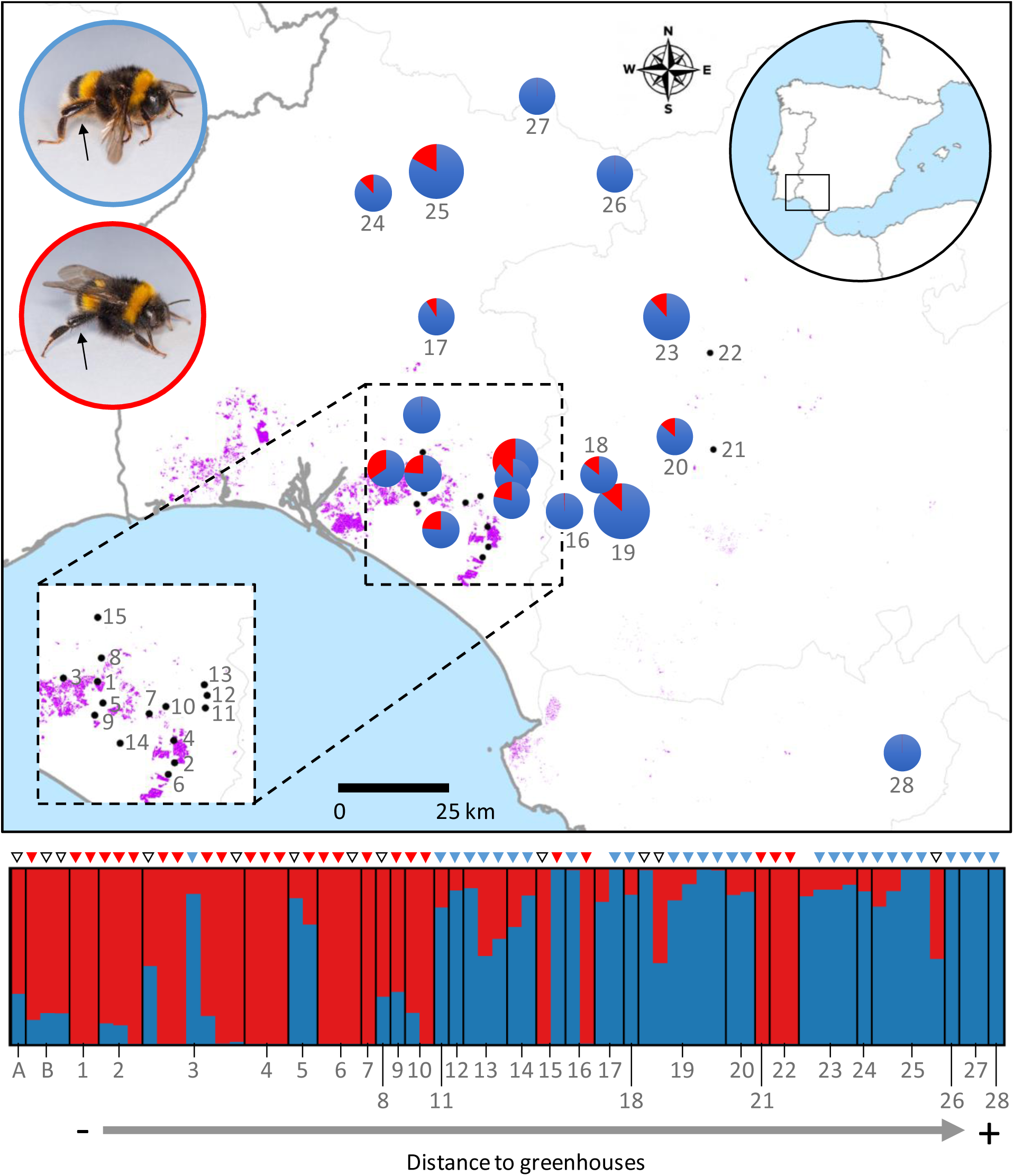
Hybridization between native and commercial bumblebees. Map showing greenhouse areas (in purple) and sampling localities (black dots and pie charts) in southwestern Spain. Pie charts (size proportional to number of genotyped individuals) show the posterior probabilities of assignment (*q*) to the native (blue) and non-native (red) genetic cluster inferred by STRUCTURE program for each sampling locality. Pie charts are only shown for localities with individuals assigned with a probability >30% to the native genetic cluster (i.e. conservatively excluding potential commercial individuals with high admixed ancestry). Bar plot shows the genetic assignment for all genotyped individuals, including four specimens sampled from commercial colonies (localities A and B) and 64 wild-caught individuals collected at 28 localities (sorted by distance to main greenhouse area). Inset pictures on top panel show a native *B. t. lusitanicus* (top, blue border) and a typical commercial individual (bottom, red border), with arrows indicating their respective reddish and black setae used to tentatively assign genotyped specimens to native, non-native and hybrid (intermediate) phenotypes (respectively blue, red and white triangles on top of STRUCTURE bar plot). Locality codes as in Table S1.

### 2.2. Phenotypic data

The main phenotypic trait characterizing the Iberian native subspecies *B. t. lusitanicus* is its distinctive legs with reddish setae, in contrast with the black or dark brown setae present in the subspecies *B. t. dalmatinus* and *B. t. terrestris*, which are most commonly used in commercial lines (Rasmont et al., 2008) (see Figure 1). We used this trait to tentatively assign sampled specimens to native, non-native and hybrid (intermediate) phenotypes. The same person (F.P.M.) phenotyped all specimens without a priori information about their respective genotype.

### 2.3. Genomic library preparation and genomic data processing

We processed genomic DNA into one genomic library using the double-digestion restriction-site associated DNA sequencing procedure (ddRAD-seq) described by Peterson et al. (2012) with minor modifications detailed in Methods S1. We sequenced the library in a single-read 150-bp lane on an Illumina HiSeq2500 platform at The Centre for Applied Genomics (SickKids, Toronto, ON, Canada) and used the different programs distributed as part of the STACKS v.1.35 pipeline (*process_radtags*, *ustacks*, *cstacks*, *sstacks* and *populations*) to assemble our sequences into *de novo* loci and call genotypes (Catchen, Hohenlohe, Bassham, Amores, & Cresko, 2013). Methods S2 and Figure S1 provide all details on sequence assembling and data filtering.

### 2.4. Genetic assignment and hybrid identification

We identified hybrid and purebred individuals in our dataset using the Bayesian Markov chain Monte Carlo clustering method implemented in the program STRUCTURE v.2.3.3 (Pritchard, Stephens, & Donnelly, 2000). We conducted 15 independent runs for each value of *K* = 1-10 using 200,000 MCMC cycles after a burn-in step of 100,000 iterations, assuming correlated allele frequencies and admixture, and without using prior population information (Hubisz, Falush, Stephens, & Pritchard, 2009). We retained the ten runs having the highest likelihood for each value of *K* and identified the number of genetic clusters best fitting the data using the Δ*K* method (Evanno, Regnaut, & Goudet, 2005). In STRUCTURE, the posterior probability (*q*) describes the proportion of an individual genotype originating from each of the *K* genetic clusters. We considered a *q*-value of 0.95 to classify individuals as purebreds or hybrids, an adequate threshold according to validation analyses based on genotypes simulated for different hybrid classes (F1, F2, and first generation backcrosses) using HYBRIDLAB (Nielsen, Bach, & Kotlicki, 2006) and STRUCTURE and using the same settings than for our empirical dataset (Vähä & Primmer 2006) (see Methods S3, Table S2 and Figure S2). Complementary to Bayesian clustering analyses and in order to visualize the major axes of genomic variation, we performed an individual-based principal components analysis (PCA) using the R v.3.3.3 (R Core Team 2019) package *adegenet* (Jombart, 2008).

## 3. RESULTS

### 3.1. Genomic dataset

We obtained 132,125,053 reads (mean ± SD = 1,887,500 ± 610,189 reads/individual) across all genotyped individuals, of which 92% were retained after the different quality filtering steps in STACKS (Figure S1). After removing one of two individuals identified as full-siblings and another individual with a very low sequencing depth, the final dataset contained 68 non-sibling individuals (see Methods S2). The final exported dataset obtained with STACKS after removing loci that did not meet the population filtering requirements retained 9,063 single SNP loci.

### 3.2. Genetic assignment and hybrid identification

Log probabilities [Pr(X|*K*)] of STRUCTURE analyses for the empirical dataset sharply increased from *K* = 1 to *K* = 2 and steadily from *K* = 2 to *K* = 10 (Figure S3). The Δ*K* method (Evanno et al., 2005) indicated that the best-supported number of clusters was *K* = 2 (Figure S3). Visual inspection of the specimens collected in the field clearly show individuals with *B. t. lusitanicus* phenotypes and individuals with commercial phenotypes or mix characteristics (Figure 1). Accordingly, one genetic cluster corresponded to the phenotypes of the native *B. t. lusitanicus* (hereafter, native genetic cluster) and the other to commercial phenotypes (hereafter, non-native genetic cluster) (Figure 1; see Section 3.3 for more details). Principal components analyses (PCA) also supported a clear separation along PC1 between the two genetic clusters identified by STRUCTURE analyses, with hybrid/introgressed individuals placed at an intermediate position (Figure 2). Accordingly, posterior probabilities of assignment (*q*) yielded by STRUCTURE were highly correlated with scores obtained for the first principal component (PC1) (Pearson’s correlation, *r* = 0.98, *P* < 0.001).

**FIGURE 2.**
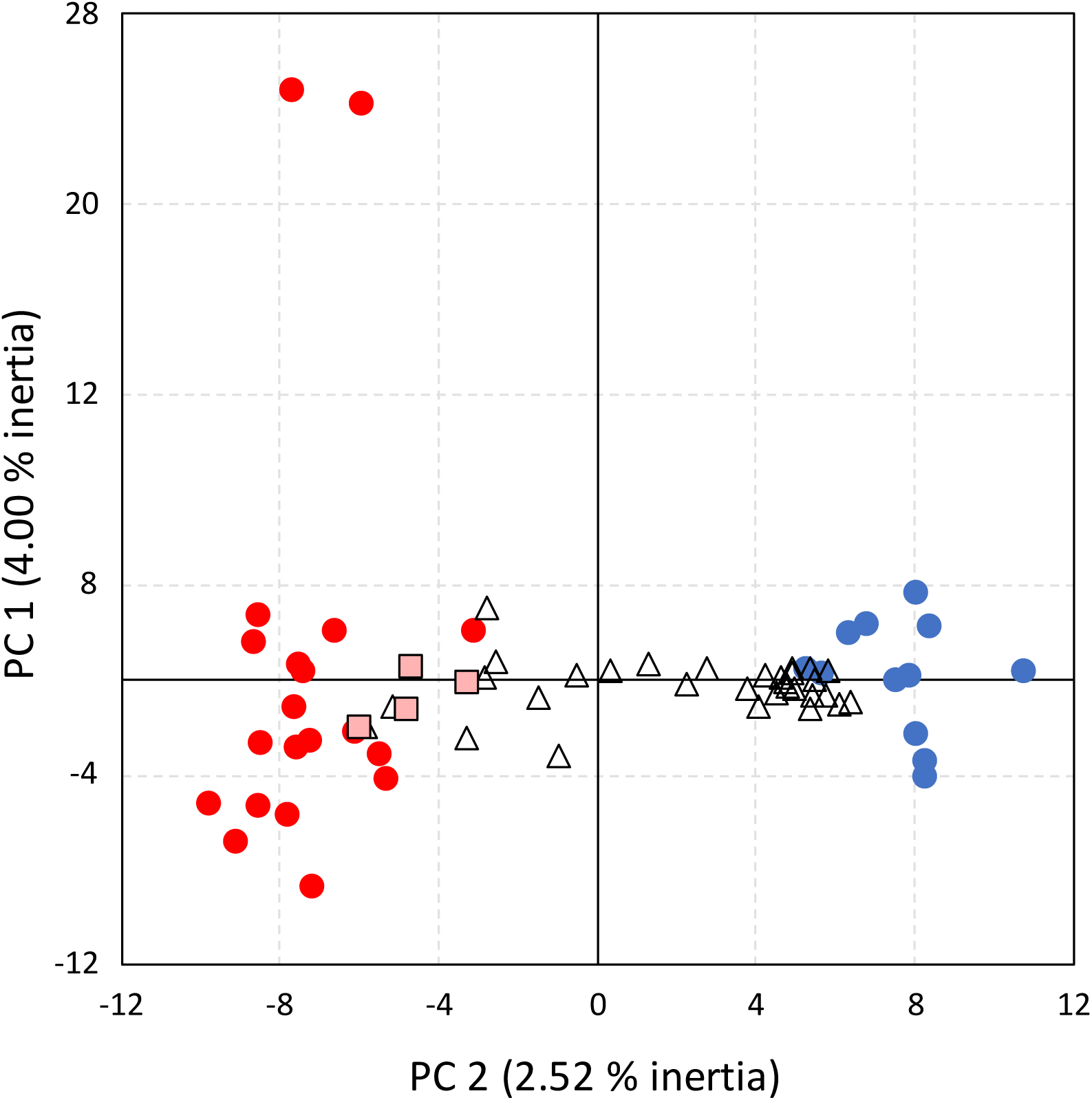
Principal component analysis (PCA) of genomic variation for 68 individuals of the bumblebee *Bombus terrestris*. Dots represent individuals assigned to the native (blue) and non-native (red) genetic clusters and open triangles indicate hybrids between them according to STRUCTURE analyses considering a threshold *q*-value of 0.95. Light red squares indicate the four specimens sampled from commercial colonies.

Considering a threshold *q*-value of 0.95 to classify genotypes as purebreds or hybrids, 14 individuals were assigned with a high probability to the native genetic cluster, 20 individuals were assigned to the non-native genetic cluster, and 30 individuals were hybrids with different levels of genetic admixture (Figure 3). Around 31% of wild-caught specimens were assigned with a high probability to the non-native genetic cluster, indicating that they represent either commercial individuals foraging in natural or semi-natural areas or the presence of colonies established in the wild from naturalized individuals. As expected, most of these individuals were sampled nearby the main greenhouse areas. Interestingly, the four individuals sampled from colonies of the two companies operating in the area were assigned with a probability of 14-29% to the native genetic cluster. This indicates that commercial lines are complex breeds probably originated from a mix of different lineages that might involve either *B. t. lusitanicus* or a third lineage genetically closer to it and for which we do not have reference genotypes. However, the most striking result is that only 19% of analysed individuals were assigned with a high confidence to the pure native genetic cluster and 50% of sampled specimens were first generation hybrids or backcrosses between native and commercial genotypes, indicating that genetic introgression is pervasive in southern Spain (Figure 3).

**FIGURE 3.**
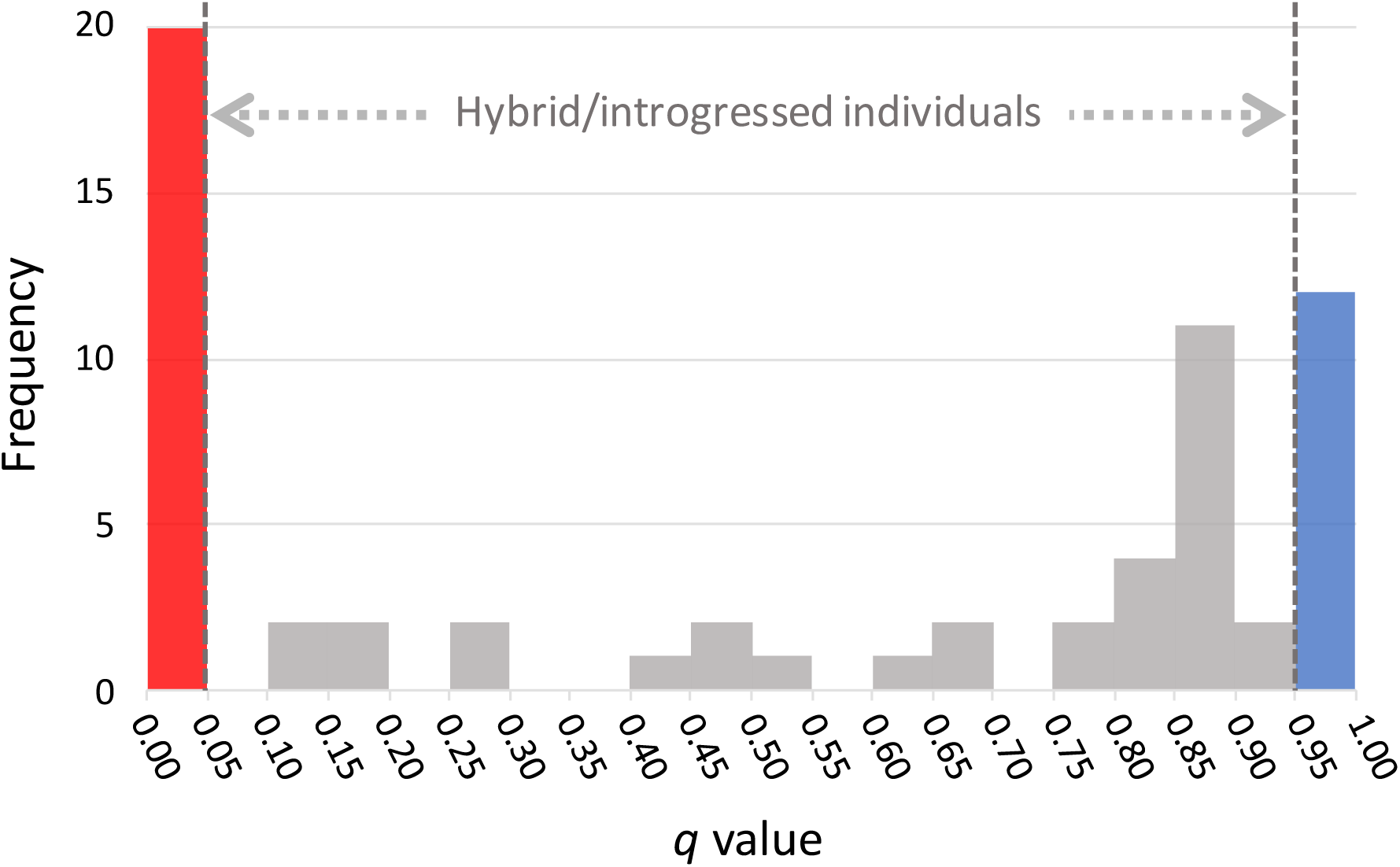
Frequency distribution of posterior probabilities of assignment (*q*-values) inferred by STRUCTURE analyses for our dataset of 64 wild-caught individuals of the bumblebee *Bombus terrestris*. Dashed vertical lines indicate threshold values of posterior probabilities (*q*-value = 0.95) used to classify individuals as purebred or hybrids. Blue and red bars indicate the frequency of purebred individuals assigned to the native and non-native genetic clusters, respectively. Grey bars indicate the frequency of individuals with different degrees of admixed ancestry (i.e. hybrid/introgressed individuals).

### 3.3. Correspondence between genotypic and phenotypic assignments

Genetic assignment scores inferred with either STRUCTURE (one-way ANOVA: *F*2,63 = 86.80, *P* < 0.001) or PCA (along PC1) (one-way ANOVA: *F*2,63 = 90.13, *P* < 0.001) were significantly different among individuals tentatively identified as natives, non-natives and hybrids based on their phenotype (*P* < 0.02 for all post hoc Tukey tests) (Figure 4). No individual assigned to the non-native genetic cluster was phenotypically identified as *B. t. lusitanicus* and 75% (9 out of 12) of individuals assigned to the native genetic cluster were tentatively classified as *B. t. lusitanicus* according to their phenotype (Figure 1). Remarkably, 63 % of the individuals tentatively identified as native based on their phenotype had some degree of introgression from the primary commercial lineage (Figure 1).

**FIGURE 4.**
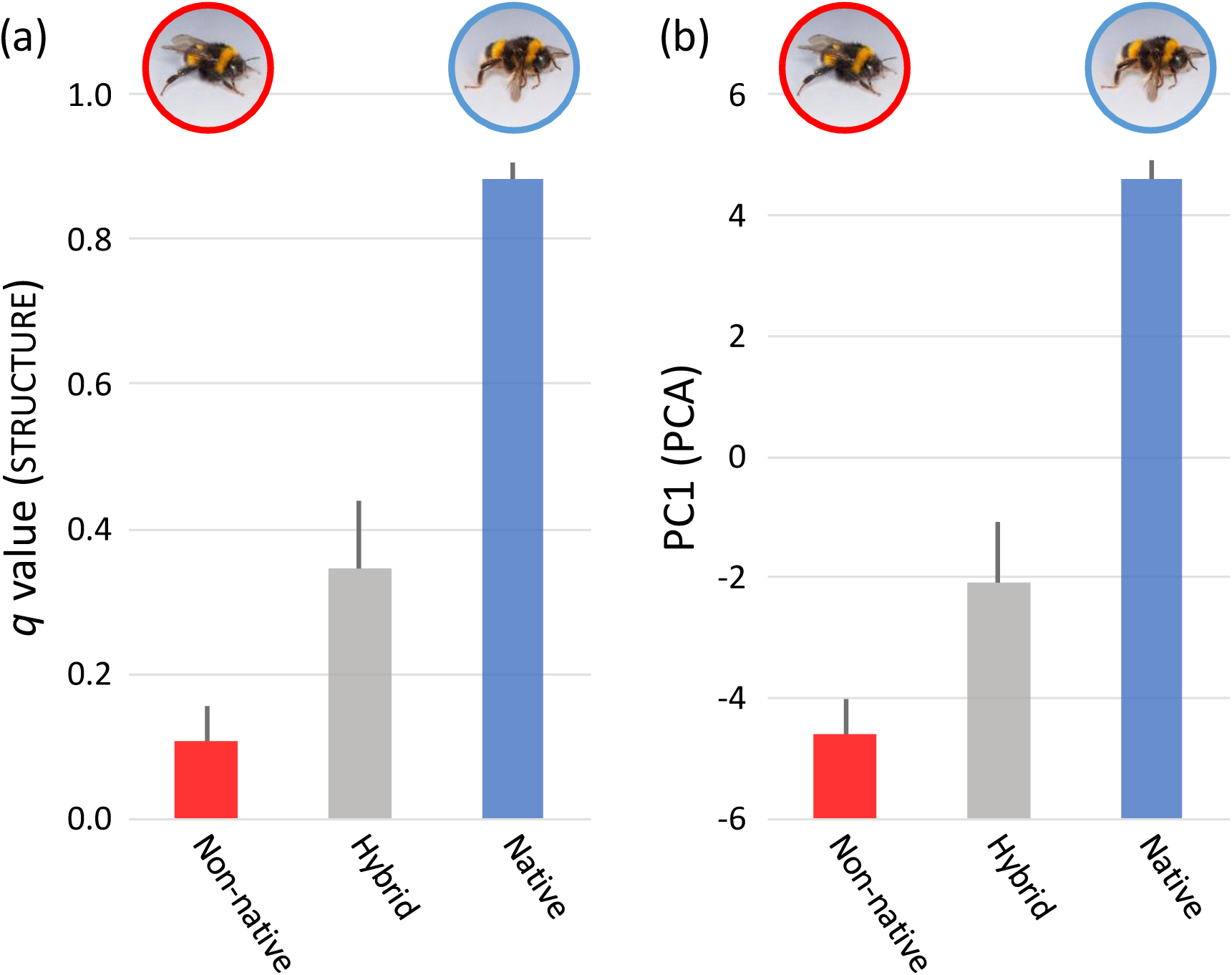
Genetic admixture scores (mean ± S.E.) obtained with (A) STRUCTURE (*q*-values) and (B) principal component analysis (PCA) (scores along PC1) for individuals that were tentatively identified as natives, non-natives and hybrids based on their phenotype.

### 3.4. Introgression spread

We found clear evidence that commercial bumblebee genotypes are spreading into native populations. The proportion of the non-native genotype decreased non-linearly with the distance to the main greenhouse areas either considering all individuals (exponential function: *F*1,26 = 35.59, *P* < 0.001, *R*2 = 0.58; Figure 5a) or only those assigned with a probably >30 % to the native lineage (i.e. conservatively excluding potential commercial individuals with admixed ancestry at a high confidence; logarithmic function: *F*1,16 = 12.86, *P* = 0.002, *R*2 = 0.45; Figure 5b). Although the frequency of commercial genotypes sharply declined with the distance to main greenhouse areas, non-native alleles have introgressed into native populations inhabiting protected natural parks >60 km away from commercial bumblebee release points.

**FIGURE 5.**
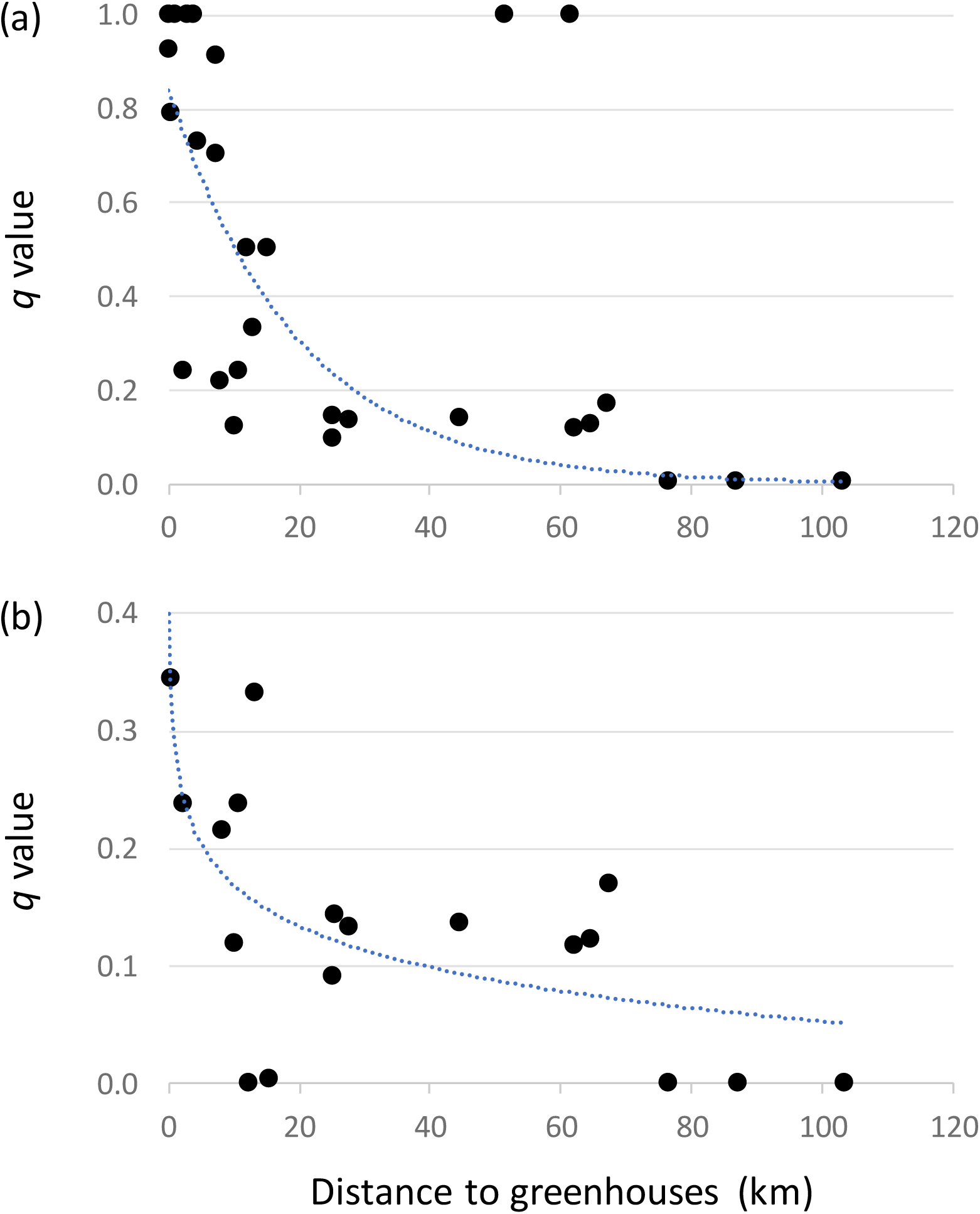
Introgression spread. Association between probability of assignment to the non-native genetic cluster and distance to main greenhouse areas considering (a) all analyzed individuals or (b) only those assigned with a probably > 30 % to the native genetic cluster (i.e. conservatively excluding with a high confidence potential commercial individuals with admixed ancestry). Regression lines represent the function yielding the highest fit to observed data (panel a, exponential; panel b, logarithmic).

## 4. DISCUSSION

The consequences of hybridization are hard to predict, but displacing the locally adapted genotype is likely to reduce the species fitness. In fact, there have been observations that native subspecies densities decline nearby greenhouses, where commercial subspecies are more prevalent (Trillo et al., 2019), indicating a potential competitive displacing. This is not surprising as the newly arrived genotypes are maintained by a huge propagule pressure, hence, their populations are subsidized and do not depend on their performance in natural conditions. As commercial lines are selected for non-random characteristics, including big colonies and fast generation times, the introgressed alleles may be suboptimal and non-locally adapted. In fact, if alleles associated with adaptation to warmer temperatures from southern Spain are lost, the effects may be deleterious in a context of populations that occur in the southern tip of the species distribution with an increasing pressure from ongoing climate warming (Kerr et al., 2015). This genetic introgression may be more widespread than initially though, but it has seldom been evaluated (but see Cejas, Ornosa, Munoz, & De la Rua, 2018; Kraus et al., 2011; Seabra et al., 2019). For example, in the UK, some colonies of the native *B. t. audax* subspecies, which historically hibernate during winter, have expanded their life cycle into the winter since early 2000’s, a date matching the introduction of commercial lines of *B. t. dalmaticus*. The possibility of genetic introgression in this context has been suggested but never evaluated.

In the past years, the regulation of commercial introductions of exotic species has advanced considerably, despite international coordination is still needed (Aizen et al., 2019). However, the regulation on the use of local subspecies is falling behind. For example, *B. terrestris* colonies are deployed in Europe without restrictions on the subspecies used with the exception of the Canary Islands (Spain), Norway, and since 2015, the UK. Imports of non-local bumblebees are also restricted in some West Asian countries such as Turkey and Israel. The case of the Canary Islands is exemplar, as only colonies of the locally native *B. t. canariensis* have been used since 1994. In contrast, the UK only recently tightened regulations to prevent the use of non-native subspecies, which were previously allowed to be purchased. In response, commercial producers supply since 2015 the UK sub-species *B. t. audax*.

While stronger regulations may be seen as a threat to farmers depending on such resources, we do not believe it will be problematic. Two examples are illustrative here. First, the Norwegian authorities do not allow the importation of colonies from outside the country and required the local production of the endemic subspecies *B. t. terrestris*. As the Norwegian market is small, the three big companies dominating the market refused to supply this service, but two local companies emerged to fill the gap. In the Middle East, the Israeli authorities do not allow the importation of bumblebee colonies from outside the country. As the Israeli market is big, it was profitable for one of the leading international companies to move production operations within the country in order to supply the local market. In both cases, the trade of colonies is minimized, with the increased benefit of avoiding the spread of novel pathogens.

As the market expands, more countries in North Africa and West Asia will demand for pollinator services. Strong regulations from day one are necessary to ensure preserving genetic diversity and local adaptations of native populations. Beyond bumblebees, the demand for the development of other commercial pollinators is increasing. Mason bees (*Osmia* spp.) are already being sold as pollinators in Europe and the US with no clear regulations on its shipment. For example, *Osmia bicornis* from Germany can be bought and released elsewhere in Europe with no restrictions. There are three distinctive subspecies of *O. bicornis* (*O. b. bicornis*, *O. b. cornigera*, and *O. b. fractinoris*) and the above exposed evidence with bumblebees teach us that extra care is needed when moving them outside their respective native ranges. A similar scenario happens in the US, where *O. lignaria* has two distinct subspecies (*O. l. propinqua* and *O. l. lignaria*) which should not be moved beyond their native ranges. As far as we know, trade of *O. lignaria* subspecies outside its distribution range is not happening, because the leading *Osmia* seller in the US (CrownBees) has regulated this on a personal decision, but no legislation is in place preventing this to happen when the market expands.

We urge countries to issue regulations enforcing the use of local genetic lines. The demand for commercial pollination services will keep increasing in parallel with the demand for pollinator-dependent crops (Aizen, Garibaldi, Cunningham, & Klein, 2008) and the technification of cropping systems around the world. As new pollinator taxa will be domesticated in the future to fulfil this demand, the time is ripe to recognize the high risks commercial pollinators entail for the genetic diversity of local native pollinators and act consequently. Breeding of local genetic lines has also the potential to minimize transport and spread of pathogens and to create opportunities in the local markets. We cannot allow commercial arguments to overrule ecological considerations.

## AUTHOR CONTRIBUTIONS

I.B. and J.O. designed the study; F.P.M. and A.H-G. conducted field and laboratory work; J.O. analysed the genomic data; I.B. drafted the manuscript and all authors contributed to writing.

## ACKNOWLEDGEMENTS

We thank to Centro de Supercomputación de Galicia (CESGA) and Doñana’s Singular Scientific-Technical Infrastructure (ICTS-RBD) for access to computational resources. Logistical support was provided by Laboratorio de Ecología Molecular from Estación Biológica de Doñana (LEM-EBD). This work is part of the project “ABEJORROS” granted by “Ayudas Fundación BBVA a equipos de investigación científica 2018”.

## DATA ACCESIBLITY

Raw Illumina reads have been deposited at the NCBI Sequence Read Archive (SRA) under BioProject (ref number will be provided upon acceptance). Input files for all analyses are available for download at FigShare (doi: available upon acceptance).

## CONFLICTS OF INTEREST

The authors declare no competing interests.

## ORCID

*Ignasi Bartomeus* https://orcid.org/0000-0001-7893-4389

*Joaquín Ortego* https://orcid.org/0000-0003-2709-429X

## SUPPORTING INFORMATION

Additional supporting information may be found online in the Supporting Information section at the end of the article.

## SUPPLEMENTAL METHODS

**METHODS S1** Genomic library preparation

We used NucleoSpin Tissue kits (Macherey-Nagel, Durën, Germany) to extract and purify genomic DNA from the hind femur of each individual. We processed genomic DNA into one genomic library using the double-digestion restriction-site associated DNA sequencing procedure (ddRAD-seq) (Peterson, Weber, Kay, Fisher, & Hoekstra, 2012). In brief, we digested DNA with the restriction enzymes *Mse*I and *Eco*RI (New England Biolabs, Ipswich, MA, USA) and ligated Illumina adaptors including unique 7-bp barcodes to the digested fragments of each individual. We pooled ligation products and size-selected them between 475-580 bp with a Pippin Prep machine (Sage Science, Beverly, MA, USA). We amplified the fragments by PCR with 12 cycles using the iProofTM High-Fidelity DNA Polymerase (BIO-RAD, Hercules, CA, USA) and sequenced the library in a single-read 150-bp lane on an Illumina HiSeq2500 platform at The Centre for Applied Genomics (SickKids, Toronto, ON, Canada).

**METHODS S2** Genomic data filtering and assembling

We used the different programs distributed as part of the STACKS v.1.35 pipeline (*process_radtags*, *ustacks*, *cstacks*, *sstacks*, and *populations*) to assemble our sequences into *de novo* loci and call genotypes (Catchen, Hohenlohe, Bassham, Amores, & Cresko, 2013; Catchen, Amores, Hohenlohe, Cresko, & Postlethwait, 2011; Hohenlohe et al., 2010). We de-multiplexed and filtered reads for overall quality using the program *process_radtags*, retaining reads with a Phred score > 10 (using a sliding window of 15%), no adaptor contamination, and that had an unambiguous barcode and restriction cut site. At this step, we removed one individual due to a very low sequencing depth (< 100,000 retained reads) (Figure S1; Table S1). For the rest of genotyped individuals, we screened raw reads for quality with FASTQC v.0.11.5 (Simon, 2018) and trimmed all sequences to 129-bp using SEQTK (Heng, 2017) in order to remove low-quality reads near the 3’ ends. We assembled filtered reads *de novo* into putative loci with the program *ustacks*. We set the minimum stack depth (*m*) to three and allowed a maximum distance of two nucleotide mismatches (M) to group reads into a “stack”. We used the “removal” (*r*) and “deleveraging” (*d*) algorithms to eliminate highly repetitive stacks and resolve over-merged loci, respectively. We identified single nucleotide polymorphisms (SNPs) at each locus and called genotypes using a multinomial-based likelihood model that accounts for sequencing errors, with the upper bound of the error rate (*ε*) set to 0.2 (*3-5*) (Catchen et al., 2013; Catchen et al., 2011; Hohenlohe et al., 2010). We built a catalogue of loci using the *cstacks* program, with loci recognized as homologous across individuals if the number of nucleotide mismatches between consensus sequences (*n*) was ≤2. We matched each individual data against this catalogue using the program *sstacks* and exported output files in different formats for subsequent analyses using the program *populations*. For all downstream analyses, we exported only the first SNP per RAD locus (option *write_single_snp*) and retained loci with a minimum stack depth ≥ 5 (*m* = 5), that were sequenced in at least 50% of the individuals of each locality (parameter *r* = 0.5), represented in ∼66% of localities (parameter *p* = 20), and with a minimum minor allele frequency (MAF) ≥ 0.01 to reduce the number of false polymorphic loci due to sequencing errors. We used the option *relatedness2* in VCFTOOLS (Danecek et al., 2011) to calculate relatedness among all pairs of genotyped individuals (Manichaikul et al., 2010) and determine the existence of full-siblings in our sample, which might happen if we have collected multiple workers from the same colony (e.g. Jackson et al., 2018). These analyses showed that only two individuals from the same locality had relatedness values compatible with a full-sibling relationship (relatedness = 0.34) and we only retained one of them (the one with the highest sequencing depth) for subsequent analyses (e.g. Jackson et al., 2018) (Table S1). The rest of the individuals had negative relatedness values (ranging from −0.01 to −21.94), which excludes the possibility of a sib relationship (Manichaikul et al., 2010). The final dataset contained 68 non-sibling individuals (Table S1).

**METHODS S3** Validation of genetic markers and sampling scheme for hybrid identification

The proper identification of hybrid and purebred individuals depends on many aspects, including the degree of genetic differentiation between parental taxa, the number of loci employed, and the overall proportion of hybrids in the sample (McFarlane & Pemberton, 2019; Vähä & Primmer, 2006). Thus, we used simulations to (i) determine the power of our set of SNP markers to infer hybridization and (ii) establish a proper *q*-value threshold for identifying hybrid and purebred individuals based on the probabilities of membership inferred from STRUCTURE analyses (e.g. Cullingham, James, Cooke, & Coltman, 2012; Hasselman et al., 2014). We used HYBRIDLAB (Nielsen, Bach, & Kotlicki, 2006) to simulate 50 genotypes for each parental class. To this end, we used as reference the allele frequencies of those individuals (i.e. 12 native and 12 non-native) having the highest probabilities of assignment to their respective genetic cluster (*q* > 0.95) based on STRUCTURE analyses on the empirical dataset (Hasselman et al., 2014; Lepais et al., 2009). In turn, we used simulated genotypes for each parental class to generate 10 simulated individuals for each of four hybrid categories: F1, F2 and backcrosses in both directions (i.e. a total of 40 hybrid individuals). On the basis of these simulated genotypes, we ran three independent STRUCTURE analyses that included all simulated hybrids (i.e. 10 from each hybrid class) and different numbers of parental genotypes (50, 25, and 10 individuals from each parental group). We analysed simulated datasets using the same settings as described above for STRUCTURE analyses on our empirical dataset. Finally, we calculated the accuracy, efficiency and overall performance of STRUCTURE clustering analyses for correctly identifying parental and hybrid individuals with our SNP panel (Vähä & Primmer, 2006). Considering a threshold *q*-value of 0.95 to classify individuals as purebreds or hybrids, all simulated genotypes were assigned to their correct class independently on the proportion of purebred reference individuals included in the dataset (i.e. accuracy, efficiency and performance = 100% in all cases) (Table S2 and Figure S2). Thus, our set of SNP markers has an extraordinary power of resolution to identify hybrid and purebred individuals (Vähä & Primmer, 2006).

## SUPPLEMENTAL TABLES

**TABLE S1.** Geographical location of the sampling localities, province, locality code, sampling year, number of collected individuals per locality (*n*), latitude and longitude.

| Locality | Province | Code | Year | *n* | Latitude | Longitude |
| --- | --- | --- | --- | --- | --- | --- |
| Agrobío S.L.<sup>1</sup> | - | A | 2017 | 1 | - | - |
| Koppert España S.L.<sup>1</sup> | - | B | 2017 | 3 | - | - |
| San Cayetano | Huelva | 1 | 2017-2018 | 2 | 37.288118 | -6.675053 |
| El Rocío<sup>2</sup> | Huelva | 2 | 2017 | 4 | 37.145372 | -6.493541 |
| Lucena del Puerto | Huelva | 3 | 2017-2018 | 7 | 37.291847 | -6.752693 |
| Polígono Industrial Matalagrana | Huelva | 4 | 2017 | 3 | 37.185900 | -6.497318 |
| Las Vaquerizas | Huelva | 5 | 2017-2018 | 2 | 37.249358 | -6.660825 |
| La Rocina | Huelva | 6 | 2017-2018 | 3 | 37.123693 | -6.506917 |
| Pinar de Jurado | Huelva | 7 | 2017 | 1 | 37.232950 | -6.555126 |
| Bonares | Huelva | 8 | 2017 | 1 | 37.331339 | -6.668110 |
| Arboreto Del Villar | Huelva | 9 | 2017 | 1 | 37.226528 | -6.678222 |
| Almonte | Huelva | 10 | 2017-2018 | 2 | 37.247133 | -6.517965 |
| Casa de Santa Teresa | Huelva | 11 | 2018 | 1 | 37.247232 | -6.428382 |
| Pinares de Hinojos | Huelva | 12 | 2017 | 1 | 37.270273 | -6.425511 |
| Dehesa de Propios<sup>3</sup> | Huelva | 13 | 2017 | 4 | 37.289442 | -6.432840 |
| Los Cabezudos | Huelva | 14 | 2017 | 2 | 37.177224 | -6.618234 |
| Niebla | Huelva | 15 | 2017 | 2 | 37.404970 | -6.680077 |
| Villamanrique de la Condesa | Sevilla | 16 | 2017 | 2 | 37.216455 | -6.303496 |
| Zalamea la Real | Huelva | 17 | 2017 | 2 | 37.603011 | -6.656046 |
| Casa de Mario | Sevilla | 18 | 2017 | 1 | 37.291260 | -6.222771 |
| Aznalcázar | Sevilla | 19 | 2017-2018 | 6 | 37.234447 | -6.169033 |
| El Carambolo | Sevilla | 20 | 2017 | 2 | 37.387067 | -6.037275 |
| Universidad Pablo de Olavide | Sevilla | 21 | 2018 | 1 | 37.357498 | -5.933129 |
| Esquivel | Sevilla | 22 | 2018 | 2 | 37.552636 | -5.949147 |
| Guillena | Sevilla | 23 | 2017 | 4 | 37.631859 | -6.071277 |
| El Manzano | Huelva | 24 | 2018 | 1 | 37.856464 | -6.829981 |
| Fuenteheridos | Huelva | 25 | 2017-2018 | 5 | 37.905035 | -6.672912 |
| Santa Olalla del Cala | Huelva | 26 | 2017 | 1 | 37.907817 | -6.207026 |
| Cabeza la Vaca | Badajoz | 27 | 2017 | 2 | 38.060060 | -6.425208 |
| Grazalema | Cádiz | 28 | 2017 | 1 | 36.756299 | -5.436198 |

<sup>1</sup> Individuals sampled from commercial colonies
<sup>2</sup> One individual from this locality had a very low sequencing depth (< 100,000 retained reads) and we excluded it from all downstream analyses (see Figure S1)
<sup>3</sup> Relatedness analyses showed that two individuals from this locality were half-siblings and we only retained one of them for subsequent analyses

**Table S2.** Accuracy, efficiency and overall performance of assignments by STRUCTURE considering simulated genotypes for the native and non-native genetic clusters and their hybrids. Each simulated individual was classified as purebred or hybrid considering a threshold *q*-value > 0.95. Analyses included 40 hybrids (ten individuals for each of the four hybrid classes simulated: F1, F2 and backcrosses in both directions) and different numbers of parental genotypes (50, 25, and 10 purebred individuals from each parental genetic cluster). Genotypes for each category were simulated using HYBRIDLAB based on the allele frequencies of purebred individuals (*q* > 0.95) identified by STRUCTURE from our empirical dataset.

| | $n$ | Accuracy | Efficiency | Performance |
| --- | --- | --- | --- | --- |
| (a) 140 individuals |  |  |  |  |
| Native genetic cluster | 50 | 1.00 | 1.00 | 1.00 |
| Non-native genetic cluster | 50 | 1.00 | 1.00 | 1.00 |
| Hybrids | 40 | 1.00 | 1.00 | 1.00 |
| (b) 90 individuals |  |  |  |  |
| Native genetic cluster | 25 | 1.00 | 1.00 | 1.00 |
| Non-native genetic cluster | 25 | 1.00 | 1.00 | 1.00 |
| Hybrids | 40 | 1.00 | 1.00 | 1.00 |
| (c) 60 individuals |  |  |  |  |
| Native genetic cluster | 10 | 1.00 | 1.00 | 1.00 |
| Non-native genetic cluster | 10 | 1.00 | 1.00 | 1.00 |
| Hybrids | 40 | 1.00 | 1.00 | 1.00 |

## SUPPLEMENTAL FIGURES

**FIGURE S1.**
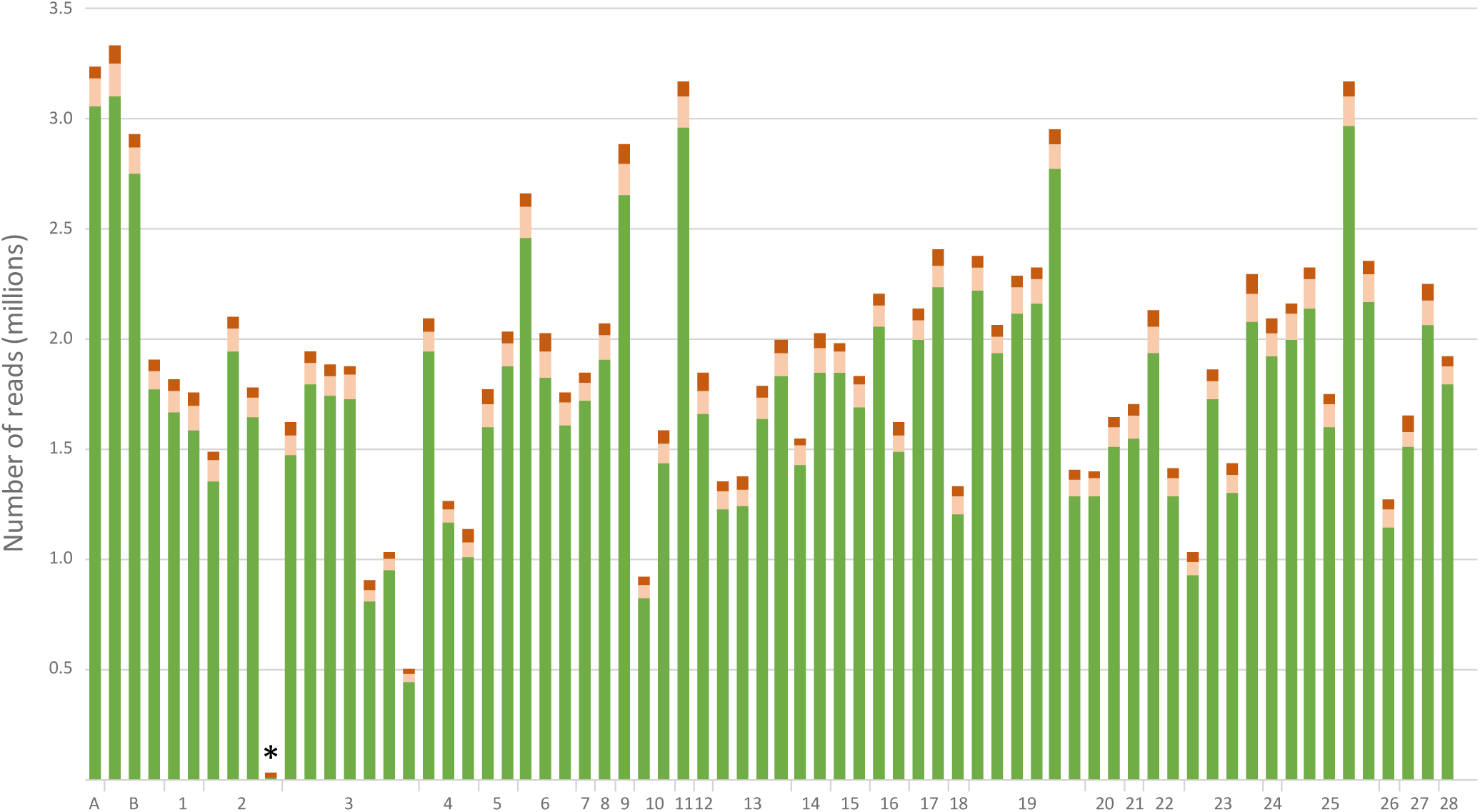
Number of reads per individual before and after different quality filtering steps by STACKS. The total height of the bars represents the total number of raw reads obtained for each individual. Within each bar, the dark red color represents the reads that were discarded by *process_radtags* due to low quality, adapter contamination or ambiguous barcode and light red color represents the reads that were discarded by *ustacks* after filtering out repetitive elements and reads that did not comply the different criteria required to create a “stack”. Green color represents the number of retained reads used to identify homologous loci. The individual marked with an asterisk had a very low sequencing depth (< 100,000 retained reads) and we excluded it from all downstream analyses. Locality codes are presented in Table S1.

**FIGURE S2.**
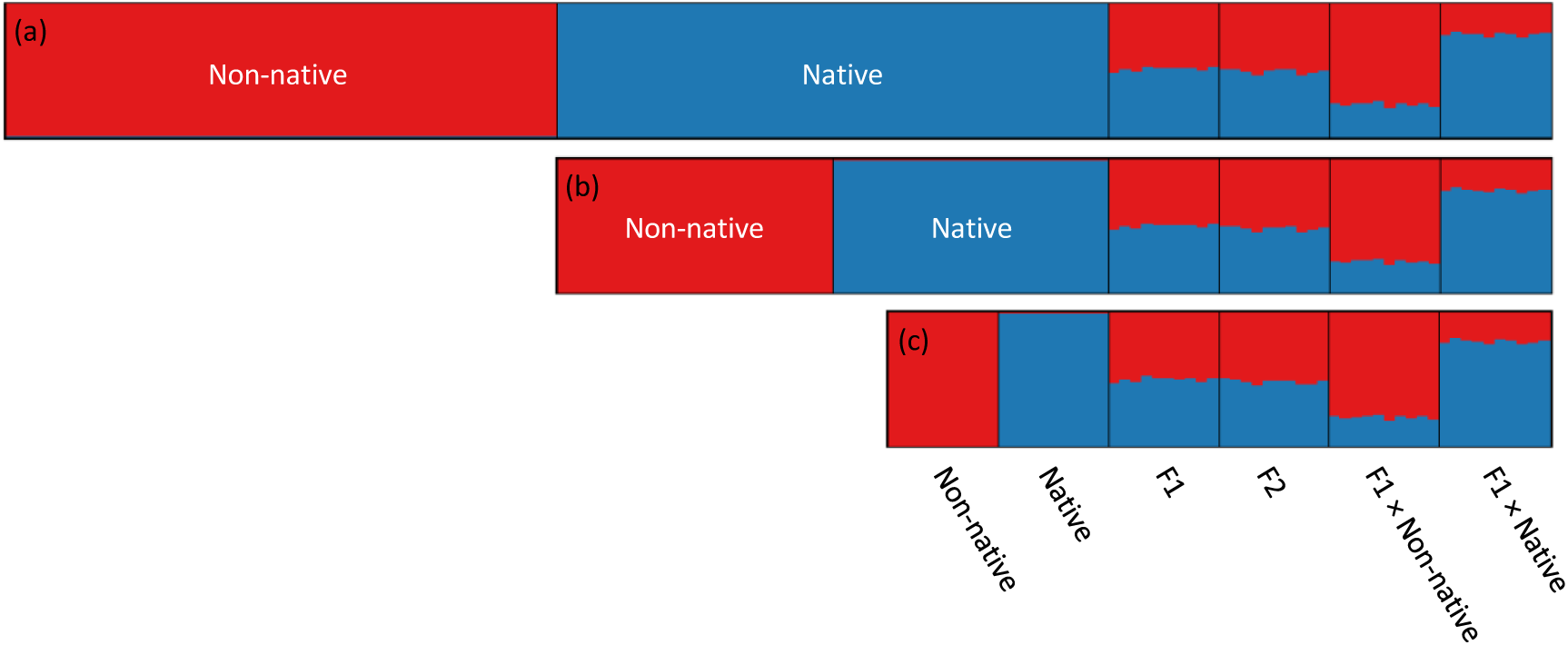
Results of assignments of simulated genotypes for the native and non-native genetic clusters and their hybrids based on the Bayesian method implemented in the program STRUCTURE. Analyses included 40 hybrids (ten individuals for each of the four hybrid classes simulated: F1, F2 and backcrosses in both directions) and different numbers of parental genotypes: (a) 50, (b) 25, and (c) 10 individuals from each parental genetic cluster. Genotypes for each category were simulated using HYBRIDLAB considering allele frequencies of purebred individuals (*q* > 0.95) identified by STRUCTURE from our empirical dataset. Each individual is represented by a vertical bar, which is partitioned into *K* colored segments showing the individual’s probability of belonging to the cluster with that color. Thin vertical black lines separate individuals from the different simulated categories.

**FIGURE S3.**
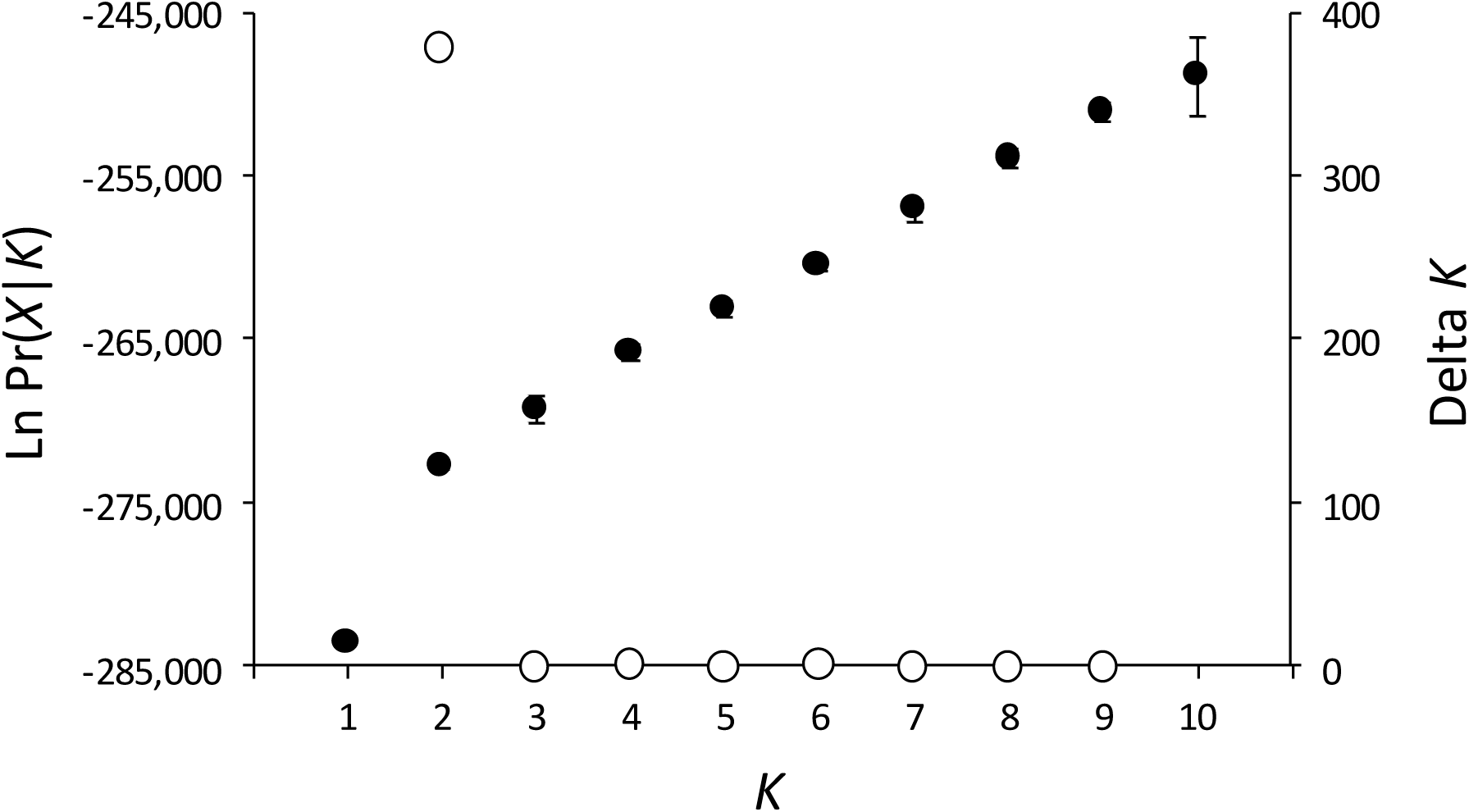
Mean (±SD) log probability of the data (Ln Pr(X|*K*)) over 10 runs of STRUCTURE (left axes, black dots and error bars) for each value of *K* and the magnitude of Δ*K* (right axes, open dots).

